# Improving the Identification Coverage of Protein Interactome by Enhancing the Click Chemistry-based Cross-linking Enrichment Efficiency

**DOI:** 10.1101/2022.01.21.475819

**Authors:** Lili Zhao, Qun Zhao, Yuxin An, Hang Gao, Xiaodan Zhang, Zhen Liang, Lihua Zhang, Yukui Zhang

**Affiliations:** CAS Key Laboratory of Separation Science for Analytical Chemistry, National Chromatographic R. & A. Center, Dalian Institute of Chemical Physics, Chinese Academy of Sciences, Dalian, Liaoning 116023, China; University of Chinese Academy of Sciences, Beijing 100039, China

**Keywords:** In vivo cross-linking, Cross-linked peptides enrichment, Click chemistry efficiency, Diverse cleavable ligands

## Abstract

Chemical cross-linking coupled with mass spectrometry has emerged as a powerful strategy which enables global profiling of protein interactome with direct interaction interfaces in complex biological systems. The alkyne-tagged enrichable cross-linkers are preferred to improve the coverage of low-abundance cross-linked peptides, combined with click chemistry for biotin conjugation to allow the cross-linked peptides enrichment. However, a systematic evaluation on the efficiency of click approaches (protein-based or peptide-based) and diverse cleavable click chemistry ligands (acid, reduction, photo) for cross-linked peptides enrichment and release is lacking. Herein, together with in vivo chemical cross-linking by alkyne-tagged cross-linker, we explored the click chemistry-based enrichment approaches on protein and peptide level with three cleavable click chemistry ligands, respectively. By comparison, the approach of protein-based click chemistry conjugation with acid-cleavable tag was demonstrated to permit the most cross-linked peptides identification. The advancement of this strategy enhanced the proteome-wide cross-linking analysis, permitting a detection of 5,017 protein-protein interactions among 1,909 proteins across all subcellular compartments with wide abundance distribution in cell. Therefore, all these results demonstrated a guideline value of our work for efficient cross-linked peptides enrichment, thus facilitated the in-depth profiling of protein interactome for functional analysis.

**GRAPHICAL ABSTRACT:** 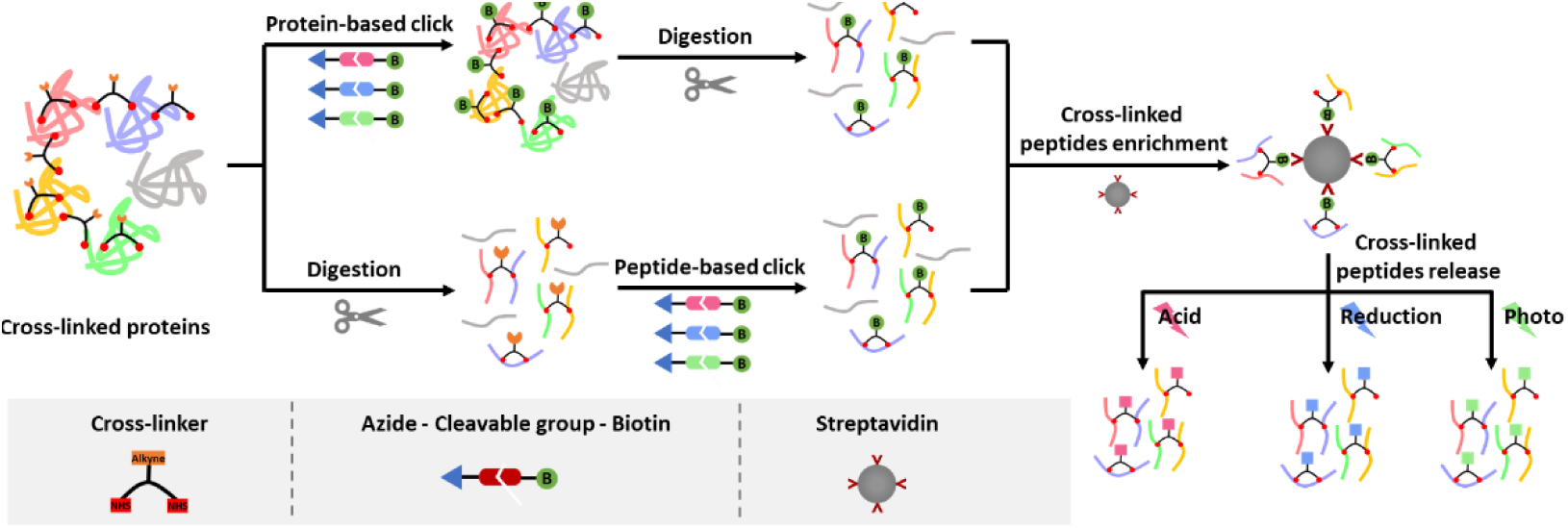

## 1. Introduction

Protein-protein interaction is one of the key regulatory mechanisms for controlling protein function and regulation in various cellular processes [1]. Nowadays, many technologies have been developed to globally study protein-protein interactions (PPIs), especially in a cellular context [2], such as affinity purification-mass spectrometry, proximity labeling techniques. Thereinto, chemical cross-linking coupled with mass spectrometry (CXMS) has recently become a powerful method for PPIs analysis with the advantage of locating the interface between interacting proteins. This strategy has been successfully employed for unraveling protein complex topology and protein-protein interacting interfaces on a proteome-wide level, especially in native cells [3-5].

In the CXMS strategy, a cross-linker is used to covalently link the active groups of amino acid residues positioned in close proximity between and within proteins. Since the cross-linkers primarily react with the amino acids on protein surface, it greatly limits the yield of cross-linking products. Exemplified by N-hydroxysuccinimidyl (NHS) ester reactive cross-linkers, which target lysine residues of the highest abundance on protein surface, multiple types of peptide mixture exist in the digested products, including regular, mono-linked, loop-linked, and cross-linked peptides. Among the peptide mixture, cross-linked peptides respect to protein interactions are the most informative species, but the least abundant [6-8]. Thus, the analysis of the low-abundance cross-linked peptides was seriously inhibited by the non-cross-linked peptides. In response, many efforts have been made to increase the relative abundance of cross-linked peptides [3, 9-12]. Among these reported methods, enrichable cross-linkers incorporated an affinity handle were the most promising. With the superiority of small steric hindrance in facilitating the cross-linker transported into the cell for in vivo cross-linking, alkyne/azide-tagged cross-linkers were increasingly used by introducing biotin with click chemistry, followed by streptavidin beads purification [3, 13, 14]. Taking advantage of this strategy, Wheat et al. identified 13,904 unique lysine-lysine linkages from in vivo cross-linked HEK 293 cells by peptide-based click chemistry, permitting construction of the largest in vivo PPI network to date [3].

Click chemistry reaction with the features of bio-orthogonality, quick reaction speed and great specificity, has been successfully applied to activity-based protein profiling, enzyme-inhibitors screening and protein labeling in proteomic analysis [15-18]. To remove biotin moiety from the enriched peptides for MS acquisition with good compatibility and deep coverage, different cleavable azide-biotin reagents, including acid-, reduction- and photo-cleavable tags were developed and used [11, 19, 20]. The current well-established protocols of click chemistry for proteomics analysis were mainly at protein level. Lately, peptide-based click chemistry was proposed for profiling the low-abundance nascent proteome with a 2-fold increase of the identified target peptides than that of protein-based click chemistry, attributed to potentially reducing steric hindrance [21]. In the study of chemical cross-linking, cross-linked peptides enrichment based on click chemistry reaction has become as a critical step for proteome-wide analysis and shows great superiority. Both protein-based [14, 22] and peptide-based [3, 13] click chemistry have been used for cross-linked peptides enrichment. However, the systematic evaluation on the effect of different click approaches and cleavable types for the cross-linked peptides enrichment and release is still unclear.

In this work, we evaluated the efficiency of alkyne-tagged cross-linker with three types of cleavable azide-biotin ligands conjugated on both protein- and peptide-based click chemistry, respectively for cross-linked peptides enrichment. The strategy presented here could provide a technological guidance for click chemistry based cross-linking enrichment, allowing in-depth PPIs analysis for charting protein interaction landscapes in cell.

## 2. Experimental section

### 2.1. Cell culture

293T cells were maintained in DMEM (Gibco, Life) and supplemented with 5% FBS (Premium, South America) and antibiotics (100 IU mL^−1^ penicillin and 100 μg mL^−1^ streptomycin) at 37 °C under 5% CO_2_ atmosphere.

### 2.2. In vivo cross-linking

The cells were harvested and washed 3 times with 1× PBS before cross-linking in centrifuge tubes. The cell pellet of 2×10^7^ cells in each group was resuspended and cross-linked in 1.2 mL 1× PBS (1% DMSO, v/v) with 5 mM BSP at room temperature for 5 min.

### 2.3. Protein-based click chemistry

(1) The cross-linked cells were collected and added 0.2% SDS (1× PBS) to extract protein. (2) Click chemistry was performed by adding cleavable azide-biotin reagent, THPTA, CuSO_4_, and sodium ascorbate to the protein sample with molar concentration ratio to the cross-linker of 1:10, 4:10; 0.5:10; 1.25:10, respectively. Volume of the reaction was 2.5 mL. The resulting mixture was rotated at 60 °C for 2 h (S-Table S1). Then, the proteins were deposited by acetone precipitation. (3) The precipitated protein pellets were air dried and resuspended in 8 M urea (50 mM NH_4_HCO_3_), following by reduction (8 mM DTT, 25 °C, 1 h) and alkylation (32 mM IAA, 25 °C, 30 min, dark), the samples were diluted to 1 M urea with 50 mM NH_4_HCO_3_ and digested with trypsin at 37 °C overnight.

### 2.4. Peptide-based click chemistry

(1) The cross-linked cells were collected and added lysis buffer (50 mM HEPES, 150 mM NaCl, pH 7.6 with 1% cocktail) to extract protein, following by reduction (8 mM DTT, 25 °C, 1 h) and alkylation (32 mM IAA, 25 °C, 30 min, dark). Then, the proteins were deposited by methanol-chloroform precipitation. The precipitated protein pellets were air dried and resuspended in 50 mM NH_4_HCO_3_, and digested with trypsin at 37 °C overnight. Next, HLB SPE cartridges was used to efficiently separate peptides from inorganic salts under neutral conditions. (2) Click chemistry was performed by adding cleavable azide-biotin reagent, TBTA, CuSO_4_, and sodium ascorbate to dried peptide digests to a final concentration of 1 mM, 1.25 mM, 10 mM, and 10 mM, respectively (S-Table S1). Volume of the reaction was 80 μL. The resulting mixture was rotated at room temperature for 2 h. Then, SCX was used for cleaning excess click chemistry reagents.

### 2.5. Cross-linked peptides enrichment

The resulting peptide mixture was incubated with streptavidin beads for 2 h at room temperature. Streptavidin-bound peptides were washed extensively before cross-linked peptides release.

### 2.6. Cross-linked peptides release

For the acid-cleavable reagent (DADPS Biotin Azide, CLICK CHEMISTRY TOOLS) ligated sample, streptavidin-bound peptides were eluted using 10% formic acid (FA) three times at room temperature. For the reduction-cleavable reagent (Azo Biotin-azide, Sigma-Aldrich) ligated sample, streptavidin-bound peptides were eluted using 300 mM Na_2_S_2_O_4_ in 20 mM HEPES, 6 M urea and 2 M thiourea buffer, pH 7.6, at room temperature. For the photo-cleavable reagent (UV Cleavable Biotin-Azide, Kerafast) ligated sample, streptavidin-bound peptides were eluted by exposing under 365 nm UV light for 1 h at room temperature. The peptides were collected, and desalted with home-made C18 Tips.

Before peptide release, the photo-cleavable reagent labeled sample should be performed with light protection, the acid-cleavable reagent labeled sample should be performed avoiding acid and not done the SCX procedure.

### 2.7. LC-MS/MS Analysis

LC-MS/MS was performed on an Orbitrap Fusion Lumos mass spectrometer (Thermo Fisher Scientific) coupled with an Easy-nLC 1200 system. A flow rate of 600 nL min^−1^ was used, where mobile phase A was 0.1% FA in H_2_O, and mobile phase B was 0.1% FA in 80% ACN and 20% H_2_O. Peptides were directly injected into the analytical column prepared in-house, with an internal diameter of 150 μm packed with ReproSil-Pur C18-AQ particles (1.9 μm, 120 Å, Dr. Maisch) to a length of approximately 30 cm. The mass spectrometry was operated in data-dependent mode with one full MS scan over the m/z range from 350 to 1500, MS scan at R = 60,000 (m/z = 200), followed by MS/MS scans at R = 15,000 (m/z = 200), RF Lens (%) = 30, with an isolation width of 1.6 m/z. MS^1^ acquisition was performed with a cycle time of 3 s. The AGC target for the MS^1^ and MS^2^ scan were 400,000 and 50,000 respectively, and the maximum injection time for MS^1^ and MS^2^ were 50 ms and 30 ms. The precursors with charge states 3 to 7 with an intensity higher than 20,000 were selected for HCD fragmentation, and the dynamic exclusion was set to 40 s. Other important parameters: default charge, 2+; collision energy, 30%.

### 2.8. Data Processing

The pLink 2 [23] software (version 2.3.9) was used for cross-links identification with precursor mass accuracy at ±10 ppm, fragment ion mass accuracy at 20 ppm, and the results were filtered by applying a 1% FDR cutoff at the spectral level. The search parameters used: instrument, HCD; precursor mass tolerance, 20 ppm; fragment mass tolerance, 20 ppm, the peptide length was set to 5-60, Carbamidomethyl[C] as fixed modification, Acetyl[Protein N-term] and Oxidation[M] as variable modification. Cross-linker was set as BSP-acid-cleave (cross-linking sites K and protein N terminus, cross-link mass-shift 376.211, mono-link mass-shift 394.222), BSP-reduction-cleave (cross-linking sites K and protein N terminus, cross-link mass-shift 411.191, mono-link mass-shift 429.201) and BSP-photo-cleave (cross-linking sites K and protein N terminus, cross-link mass-shift 390.190, mono-link mass-shift 408.201). Trypsin as the protease with a maximum of three missed cleavages allowed. The database on *Homo sapiens* was downloaded from UniProt on 2019-04-04.

## 3. Results and discussion

### 3.1. Enrichment of cross-linked peptides with different click chemistry approaches

Increasing the coverage of chemical cross-linking remains a great challenge due to the low-abundance of cross-linked peptides. As shown in Fig. 1, in vivo cross-linking was performed with our previously developed cross-linker BSP [24], which is membrane permeable, consisting of homobifunctional NHS ester reactive group and alkyne enrichable tag for living cell cross-linking in minutes. The small size and specific reactivity made alkyne group a preferred tag by introducing biotin via click chemistry for cross-linked peptides enrichment. Considering the hydrophobicity of biotin interfering with LC separation, three azide-biotin ligands of acid-, reduction- and photo-cleavable specificity with diverse solubility and size were investigated (S-Fig. S1). The acid-cleavable ligand has the longest chain and best hydrophilicity, but the whole process needs to avoid acids. The reduction-cleavable ligand has a bright yellow color from the azobenzene group, thus the binding between cross-links and the ligands could be visualized, while turned to a white color after cleavage. But the reduction-cleavable reaction occurs in a high concentration of salt, thus the desalting step is necessary. Besides, the inconvenient dark environment needs to be kept during the whole process for photo-cleavable ligand. For the acid- and photo-cleavable ligands generated sample, desalting step is no need to do in theory, which are more MS-compatible and less loss. In addition, considering the lower sample complexity and larger steric hindrance at protein-based than peptide-based click chemistry, it is necessary to investigate the effects of the three cleavable click chemistry ligands for cross-links enrichment and release, respectively.

**Fig. 1.**
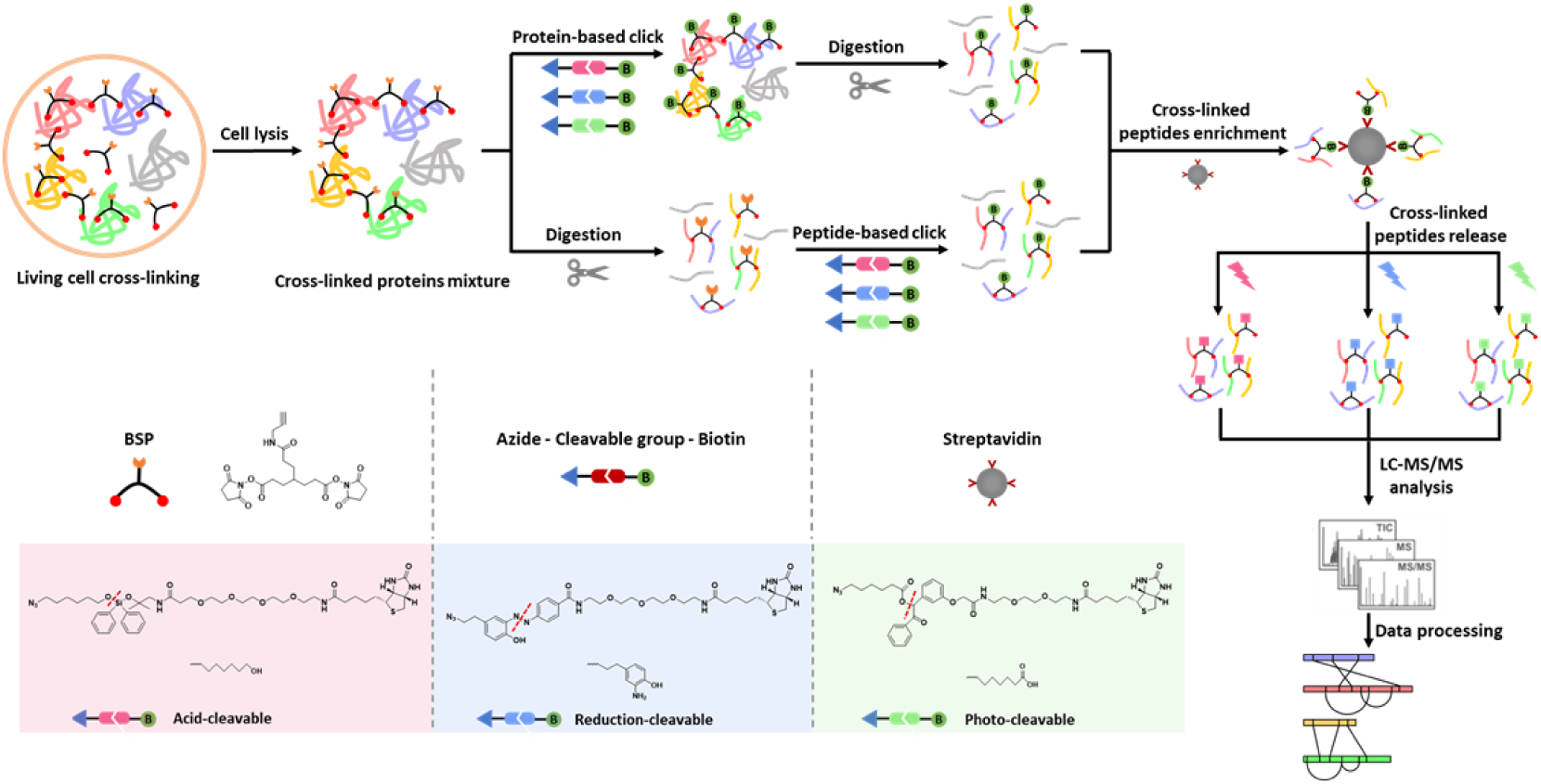
Flow diagrams of both protein-based and peptide-based click chemistry reactions for in vivo cross-links analysis. Bottom left: chemical structures of the cross-linker BSP, three cleavable azide-biotin reagents and the tags modified to the cross-linker after cleavage are respectively illustrated, in which red dashed lines represent the cleavable sites.

### 3.2. Evaluation of the cross-linked peptides enrichment efficiency

To evaluate the cross-linked peptides enrichment efficiency of the approaches with three cleavable ligands on protein and peptide-based click chemistry, we compared the type, identification number, hydrophobicity, charge character and missed cleavages of the identified cross-linked peptides. Firstly, the obviously higher proportion of cross-linked peptides was identified using the protein-based click chemistry than that with peptide-based method (Fig. 2A). This might be due to the relatively bigger steric hindrance and higher sample complexity of peptide level click chemistry reaction for cross-linked peptides. Among the six conditions, protein-based click chemistry conjugation with acid-cleavable tag identified the most cross-linked peptides, followed by peptide-based click chemistry conjugation with acid-cleavable tag. It might be attributed to the flexible spacer arm, good hydrophilicity and small reaction steric hindrance of the acid-cleavable azide-biotin reagent. For the identification with peptide-based click chemistry, 52%, 69% and 71% of the cross-linked peptides, cross-linked proteins and PPIs were commonly identified by the protein-based click chemistry method, respectively (S-Fig. S2). For the photo-cleavable tag labeled sample, protein-based click chemistry approach identified less cross-linked peptides than peptide-based approach, it might be due to the inconvenient experimental operation in dark for a longer time. While peptide-based click chemistry conjugation with reduction-cleavable tag identified the least cross-linked peptides, which might be due to the larger local hydrophobicity and steric resistance of azide, and this result was not discussed in the subsequent comparisons (Fig. 2B).

**Fig. 2.**
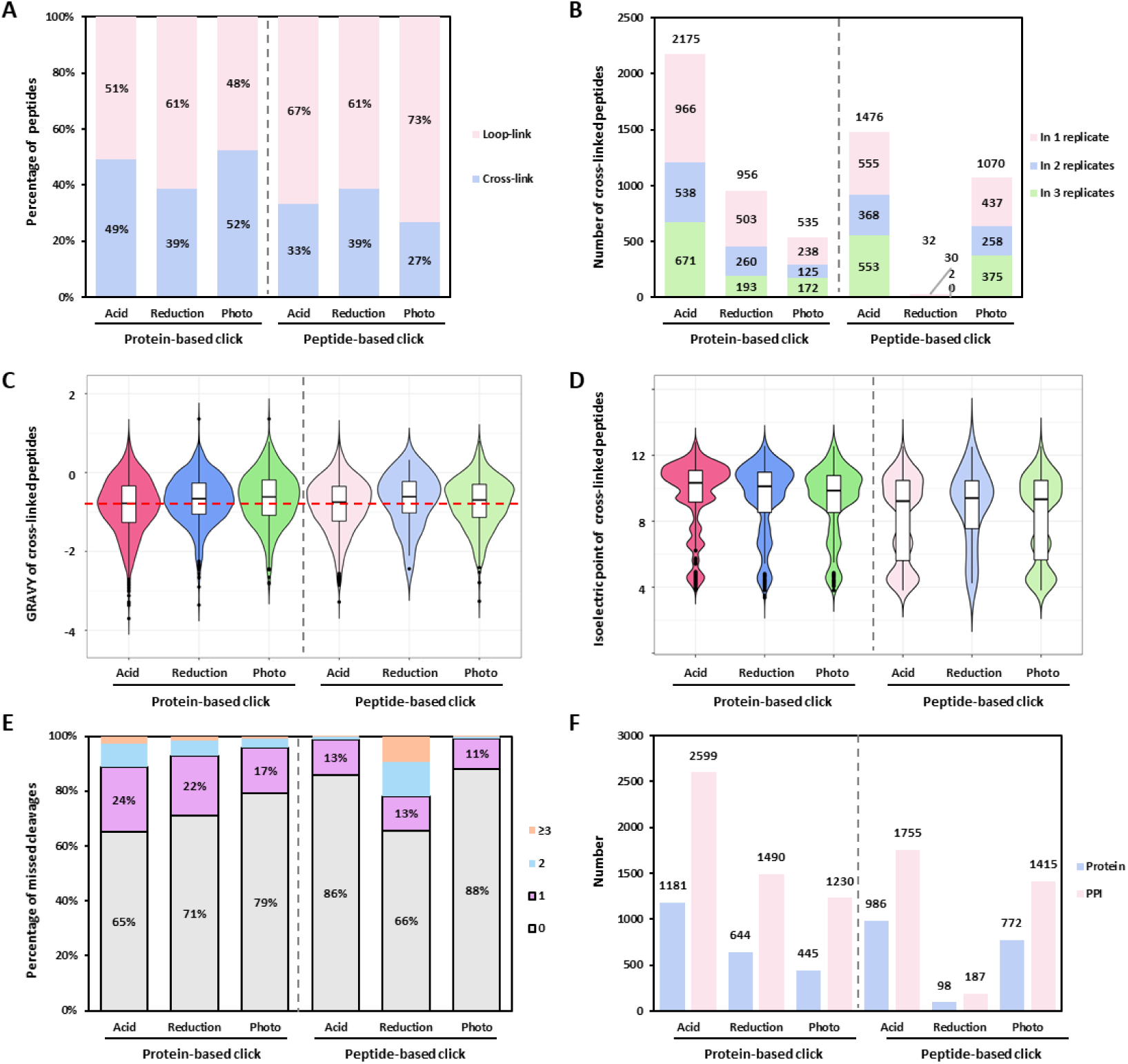
Comparison of (A) type proportion, (B) number, (C) GRAVY distribution, (D) isoelectric point distribution, (E) missed cleavage of the cross-linked peptides, as well as (F) number of the cross-linked proteins and PPIs.

Furthermore, the hydrophobic and hydrophilic performance and electrical charge of the cross-linked peptides could influence chromatographic separation and MS identification. The GRAVY distribution of the cross-linked peptides indicated higher hydrophilicity obtained for acid-cleavable tag released cross-linked peptides (Fig. 2C). The isoelectric point distribution of the cross-linked peptides demonstrated higher pI tendency on protein-based than that of peptide-based click chemistry (Fig. 2D). In addition, the missed cleavage of our identified cross-linked peptides on protein-based click chemistry was a little higher than that with peptide-based click chemistry (Fig. 2E), implying obvious steric hindrance from biotin conjugation to trypsin digestion, which could mediate pI of the peptides. Moreover, the abundance of the identified cross-linked proteins on peptide-based click chemistry was a little lower than protein-based click chemistry, but without significant difference probably attributed by the coverage of our method (S-Fig. S3). Taken together, the protein-based click chemistry conjugated with acid-cleavable azide-biotin ligand approach, which identified the most cross-linked proteins and PPIs (Fig. 2F), was recommended for cross-linked peptides enrichment to realize in-depth protein interactome identification. The detailed identification results were provided in Supporting Information 2.

### 3.3. In-depth profiling of human cell cross-linking protein interactome

To enhance the identification coverage of cross-linking, the cross-linked peptides generated from protein-based click chemistry conjugation with acid-cleavable tag were further fractionated by high-pH RPLC and identified by low-pH nanoRPLC-ESI-MS/MS analysis. Totally, 26,206 inter-protein linkages of 5,017 PPIs involved in 1,909 proteins were identified. To visualize the correlation of the identified PPIs, the interactome network was profiled, and classified according to the protein function into 173 subgroups, mainly associated with transcription and translation, protein binding, signal transduction and cell metabolism (Fig. 3A). The detailed interaction proteins classification was provided in Supporting Information 3. Cellular component analysis showed that the interacting proteins were widely distributed throughout the cell, including the plasma membrane, cytosol, endoplasmic reticulum, mitochondria, and nucleus (S-Fig. S4). In addition, we estimated the abundance range of the XL-PPI proteome. Our identified cross-linked proteins were with different abundances spanning 7 orders of magnitude. The abundance distributions of XL-PPI and MS proteomes [25] were similar, with little shift toward higher abundance proteins (S-Fig. S5), due to protein concentrations dependent of cross-linking reaction. Notably, in-depth CXMS identifications among different subcellular compartments with wide abundance distribution were greatly demonstrated in our dataset (Fig. 3B). Besides, by matching with the existing PPI databases, 69% complementarity (3,447 PPIs) was observed (S-Fig. S6), which was most likely attributed to the capture bias and distinct filter threshold of these PPI profiling methods, as well as cell type and cell state heterogeneity. These complementally identified PPIs related proteins were mainly involved in transcription and translation process, which were commonly difficult to resolve with in vitro experiment (S-Fig. S7). Furthermore, we investigated the accuracy of our obtained PPIs on account of the STRING database. Among the 5,017 PPIs, 1,456 were covered in the STRING database and matched to the corresponding reliability score. Among them, 50% (726 PPIs) of our identified PPIs were in the highest score range of 0.9-1.0, while more than 72% of the whole human PPIs data collected in the STRING database were in the relatively low score range of 0.1-0.3 (Fig. 3C), indicating the high credibility of protein interactome achieved by our method. The detailed identification results were provided in Supporting Information 4.

**Fig. 3.**
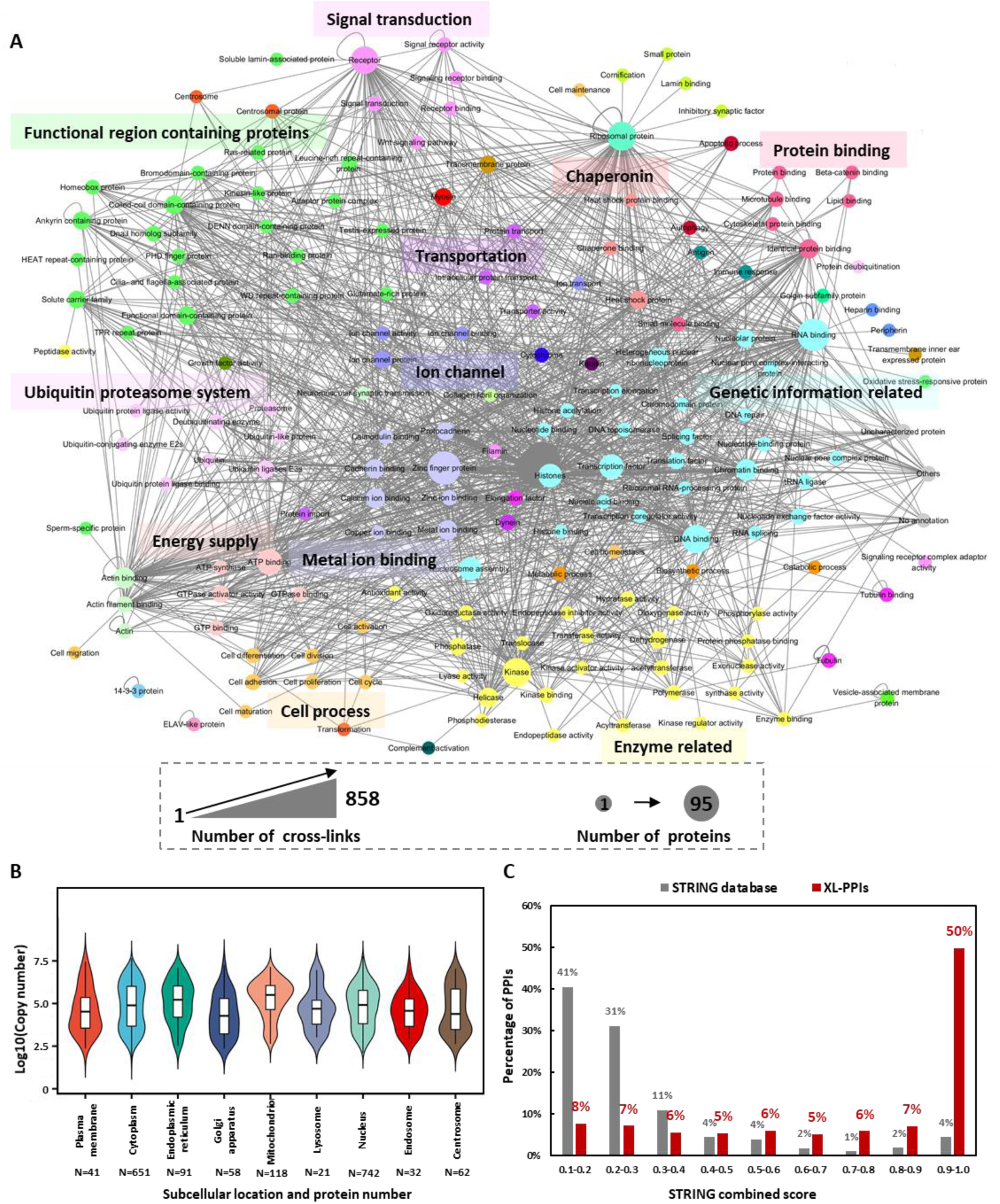
Protein-protein interaction analysis. (A) Protein interaction network mapped by our identified cross-linked proteins. The nodes are color coded based on the protein functions. Size of the nodes is proportional to the number of proteins included in the corresponding classification. Thickness of the lines represents the number of protein interactions. (B) Copy number distribution of the interaction proteins among different subcellular locations. (C) Comparison of STRING score distribution between our identified PPIs and all the human PPIs in STRING database.

## 4. Conclusions

A systematic study of the effects of different click chemistry approaches and cleavable ligands on the cross-linked peptides enrichment and identification was performed. The strategy of protein-based click chemistry conjugation with acid-cleavable ligand was proven to generate the most cross-linked peptides, while the peptide-based click chemistry conjugation with reduction-cleavable ligand identified the least. With the advancement of this strategy, the efficiency of click chemistry-based cross-linking enrichment was enhanced, and an in-depth profiling of human protein interactome with 26,206 inter-protein linkages of 5,017 PPIs involved in 1,909 proteins was constructed with in vivo cross-linking. Therefore, all these results demonstrated the great promise of our work for proteome-wide mapping of protein interaction landscapes in cell.

## CRediT author statement

Lili Zhao: Conceptualization, Methodology, Investigation, Writing Original Draft. Qun Zhao: Conceptualization, Methodology, Writing Review & Editing. Yuxin An: Methodology. Hang Gao: Provision of reagents. Xiaodan Zhang: Provision of materials. Zhen Liang: Project administration. Lihua Zhang: Conceptualization, Supervision, Project administration. Yukui Zhang: Supervision.

## Acknowledgments

This work was supported by the R&D Program of China (2018YFA0507703), National Natural Science Foundation (22074139, 21991083, 32088101, 21775150), CAS Key Project in Frontier Science (QYZDY-SSW-SLH017) and the Youth Innovation Promotion Association, CAS (2020184).

