## Supporting information1 for "Improving the Identification Coverage of Protein Interactome by Enhancing the Click Chemistry-based Cross-linking Enrichment Efficiency"

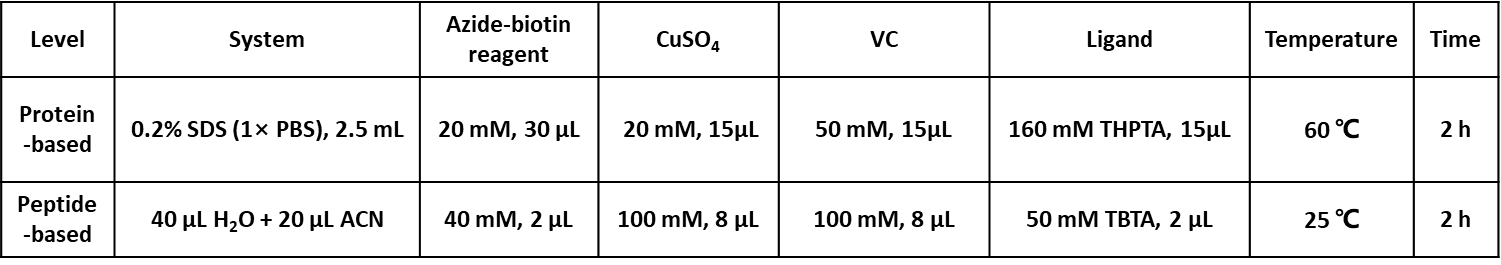

**Table S1.** Click chemistry reaction conditions.

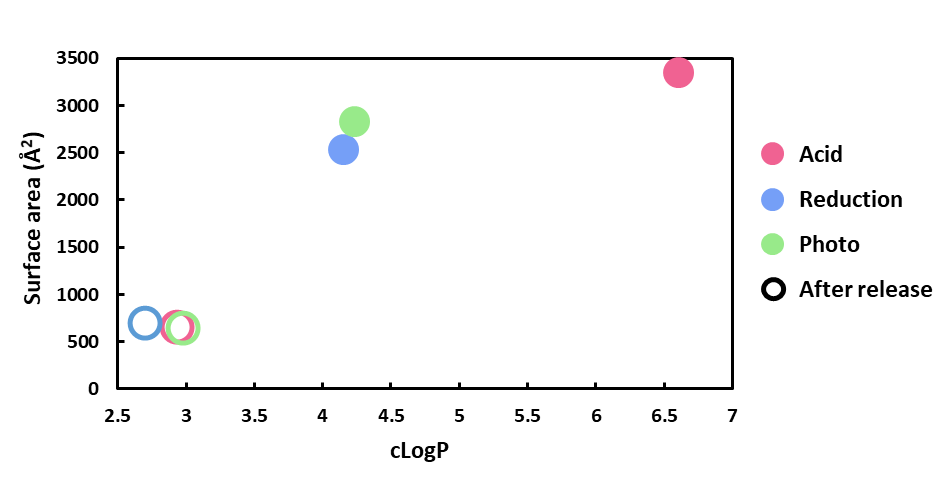

**Fig. S1.** Basic properties of the three cleavable azide-biotin reagents.

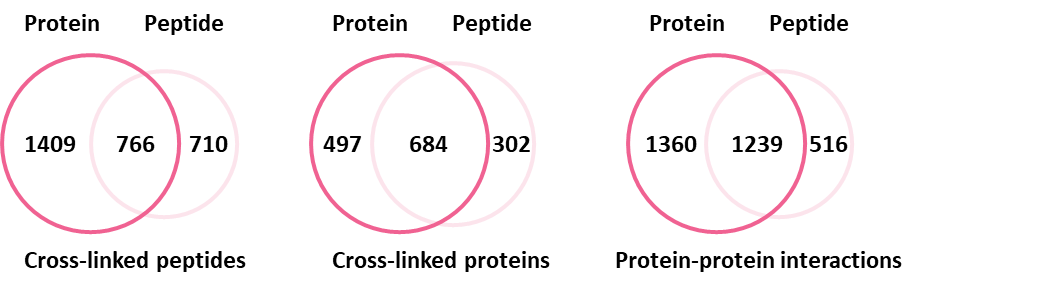

**Fig. S2.** Venn diagram of the cross-linked peptides, cross-linked proteins and PPIs from the approaches of protein and peptide-based click chemistry conjugated with acid-cleavable ligand.

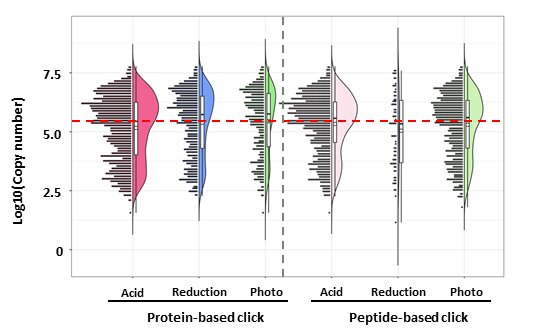

**Fig. S3.** Copy number distribution of the cross-linked proteins.

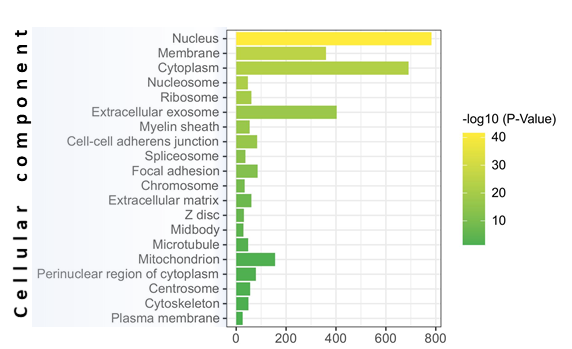

**Fig. S4.** Cellular component analysis of the interaction proteins.

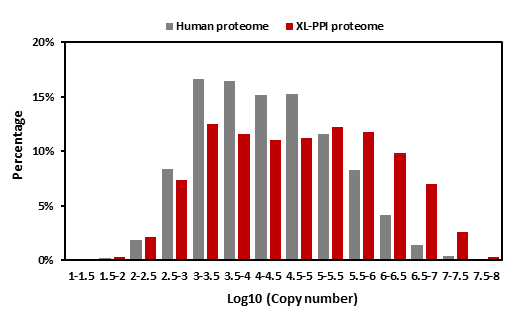

**Fig. S5.** Protein copy number distribution of our XL interaction proteome and the MS proteome of human cells.

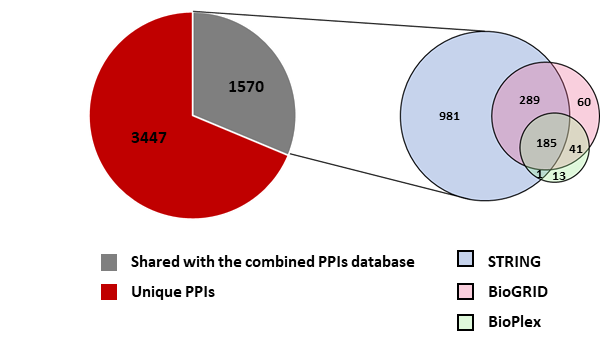

**Fig. S6.** Overlap of our identified PPIs with the existing PPIs databases of STRING, BioGRID and BioPlex.

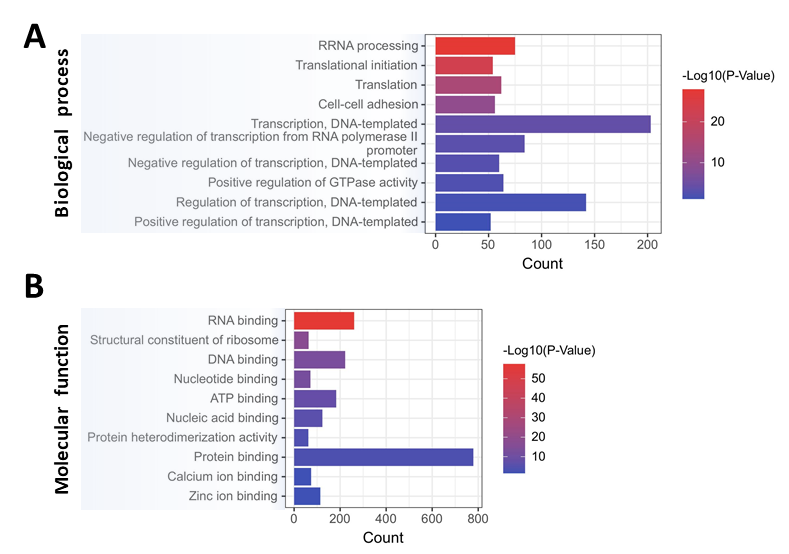

**Fig. S7.** Biological process (A) and Molecular function (B) of the existing PPI databases not reported PPIs related proteins.
